# Risk factors contributing to bacteraemia at a tertiary cancer center in South Asia

**DOI:** 10.1101/386193

## Abstract

**Context:** Cancer patients are immunocompromised due to their medical condition resulting in neutropenia, increased exposure to intravascular devices (IVDs) and prolonged hospital stays. These conditions are established risk factors in causing bacteraemia. Bateraemia is a contributing factor towards increased rates of morbidity and mortality in several countries including Sri Lanka.

**Aims:** The current study evaluates the risk factors such as demographic factors, neutrophil counts, presence of an IVD and length of hospitalization that would contribute to the development of bacteraemia among cancer patients at the Apeksha Hospital – Maharagama, Sri Lanka.

**Results:** A higher prevalence of bacteraemia compared to other countries (13.7%) was reported with the highest frequency identified from oncology wards. Patients above 60 years with carcinomas were revealed to be more susceptible. A length of hospital stay exceeding three days was a statistically significant factor in causing bacteraemia. Gram-negative organisms accounted for majority of the infections while *Acinetobacter* species were more frequently isolated from IVDs.

**Conclusions:** It could be suggested that additional care and sterility measures be taken when carrying out invasive procedures in such patients. Precautions could be taken in managing patients with a hospital stay exceeding 3 days as they have been identified as a risk group in acquiring nosocomial infections

## Introduction

Bacteraemia or bloodstream infections (BSIs) being one of the prominent cause for complications in cancer patients, has become a contributing factor towards increased rates of morbidity and mortality worldwide [1,2]. It has been identified as one of the top seven causes of death in Europe and North America [1].

The present study focused on cancer patients who were generally immunocompromised due to the underlying medical condition or treatment which includes surgery, chemotherapy and radiotherapy depending upon the type and severity or stage of cancer [3]. Previous studies had identified three major risk factors contributed to bacteraemia such as neutropenia [4,5], intravascular catheterization [6,7] and prolonged hospital stays [8,9].

Hence this study focused on evaluating the effect of three risk factors; neutropenia, intravascular catheterization and length of hospital stay towards contributing to the development of bacteraemia among cancer patients admitted to a tertiary care hospital in Sri Lanka.

## Subjects and Methods

Ethical approval for the study was obtained from the General Sir John Kotelawala Defence University Sri Lanka (KDU) Ethical Review Committee and the National Cancer Institute – Maharagama, Sri Lanka. Written consent was obtained from all participants. When recruiting children below 18 years of age, written consent was taken from the parent or guardian of child. This cross-sectional cohort study was conducted at a tertiary cancer care centre in Sri Lanka which treats only cancer patients. Blood samples were sent to the laboratory when sepsis was suspected as determined by the clinician, which included symptoms such as fever, chills and fatigue.

### Inclusion and Exclusion Criteria

All samples sent to the microbiology laboratory for blood culture, were retrospectively analyzed. Data was collected from 210 incidences during a two month period from 01st of July to the 31st of August, 2016. (Incidences correspond to the number of request forms that arrived at the laboratory during the study period)

Blood culture bottles which were refrigerated, subjected to leakage, broken, unlabelled or received without patient information and history were excluded from the study.

### Data Collection

Patients’ age, gender, hospital ward, usage of catheters, oncological diagnosis, the total white blood cell counts and neutrophil counts gathered from patients’ reports and date of admission were recorded from the respective Bed Head Tickets.

### Sample Collection and Processing

Two blood samples were collected into Blood culture bottles (BD Bactec Plus aerobic/F; Bactec Myco/F Lytic blood culture bottles and BD Bactec Ped Plus/F bottles-BD Diagnostic Systems 442192, USA) from patients with in-situ catheters; the first one through the catheter line and the second sample from a peripheral site collected at the same time [10]. All blood cultures were analyzed by the BD Bactec 9120 Automated Blood Culture System. Vials which did not provide positive results within 5 days were considered as negative for organisms [11]. Blood culture vials that were detected positive for organisms within 5 days were cultured on Blood agar, MacConkey agar and Chocolate agar [10] and incubated at 35°C overnight. The Chocolate agar plates were incubated in 5% to 10% carbon dioxide. Organism identification was performed according to routine laboratory protocols stated in Sri Lankan College of Microbiologists’ manual [10]. Clinical and Laboratory Standards Institute (CLSI) protocols were followed when performing antibiotic sensitivity testing [10].

### Determining Source of Bacteraemia

The source of bacteraemia was determined as per standard protocols stated in the Sri Lankan College of Microbiologists’ manual [10]. If blood cultures taken through the catheter becomes positive two or more hours prior to the peripheral blood culture with the same organism, it was reported as intravascular catheter associated blood stream infection (CRBSI). If the blood culture through the line was positive but the peripheral blood was negative the source of bacteraemia was determined as intravascular catheter colonization and if both the blood cultures through the catheter and the peripheral blood becomes positive with the same organism but the time gap is less than two hours or the peripheral blood culture becomes positive first, it was reported as bacteraemia not associated with intravascular catheters.

### Determining Neutropenia

Absolute Neutrophils Counts (ANCs) less than 500 cells/μl were considered as neutropenia and ANCs more than 500 cells/μl were considered as non-neutropenia [4,5].

### Determining Length of Hospital Stay

The length of hospital stay (LOS) was taken as the duration between the date of admission and the day the blood culture was detected as positive. Infections that occurred after 72 hours of hospital admittance were considered as nosocomial infections in accordance with the studies conducted by Weinstein et al. [12] and Laupland and Church [13].

### Data Entry and Statistical Analysis

Statistical Package for Social Sciences (IBM^®^ SPSS^®^ version 16.0) software was used in interpreting statistical data. The incidence of bacteraemia was analysed in terms of frequency. Two-way frequency table, cross tabulation and Chi-squared test and the odds ratio were carried out to identify whether neutropenia, intravascular catheterization and LOS contributes to the development of bacteraemia. A 95% confidence interval (95% CI) was used in all statistical analysis.

## Results

### Study Population and Demographic Characteristics

The current study identified 120 positive bacteraemic cases among 210 cancer patients over a period of 2 months from 1^st^ July 2016 to 31^st^ August 2016. The bacteraemic population of the current study consisted of 74 (35.2%) males and 46 (21.9%) females. Age was categorized based on the United Nations Provisional Guidelines, 1982 [14]. Accordingly, age groups were included in the study were Infants (0-1 year), Pre-school (2-5 years), Schooling (6-19), Working (20-60 years) and Seniors (>60 years). The overall study population consisted of 104 (49.5%) paediatric patients of which 53 (25.2%) had bacteraemia and a total of 106 (50.5%) adult patients of which 67 (31.9%) had bacteraemia. A markedly high prevalence of BSIs was recorded from the Senior category which consisted of patients aged above 60 years (Table 01). Though age was not a statistically significant factor in causing bacteraemia within a confidence interval of 95% (*P*= 0.053), it was found to be a significant causative factor at 90% CI.

**Table 01:**
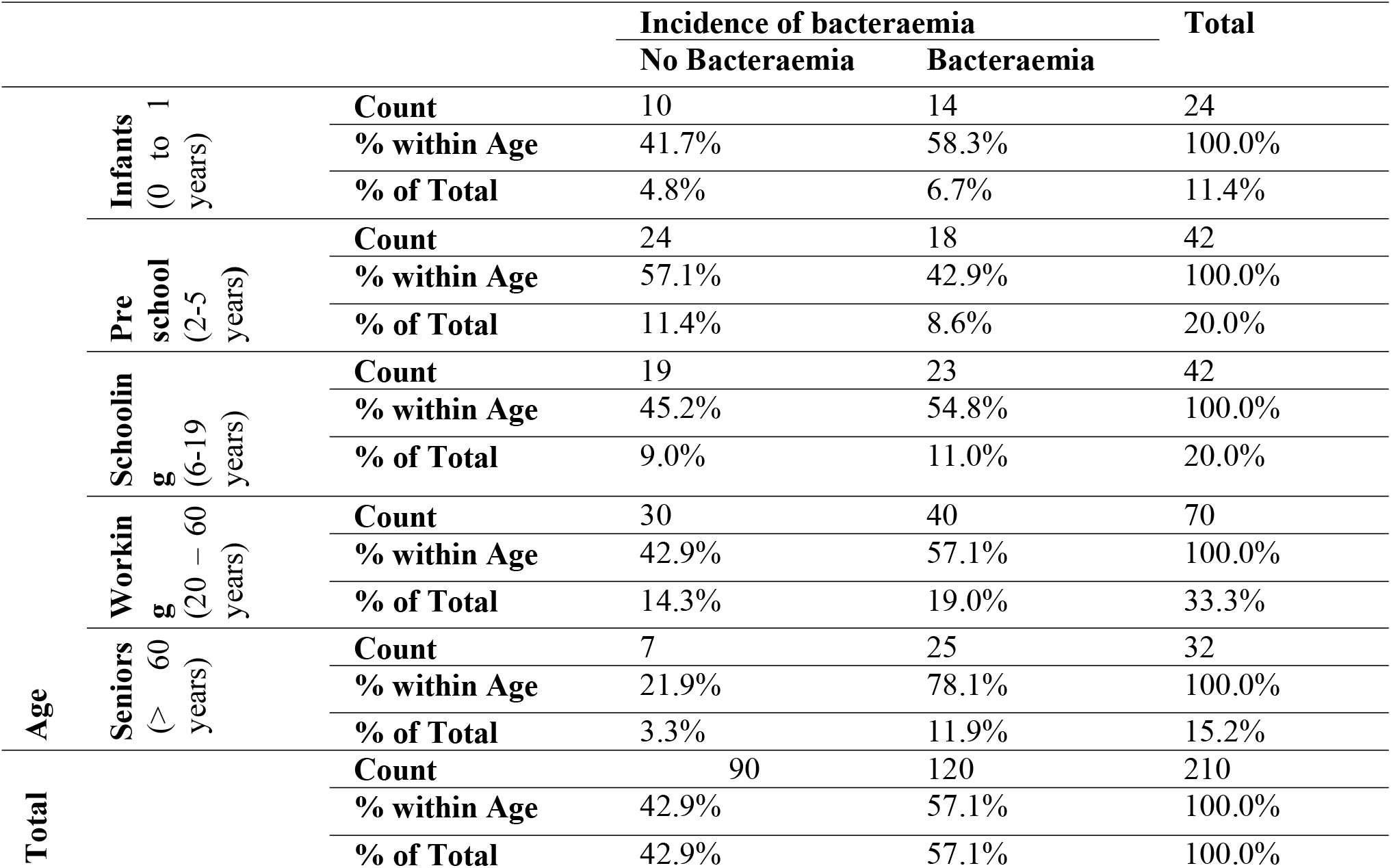
The distribution of bacteraemia within the age groups categorized according to the United Nations Provisional Guidelines, 1982

### Hospital wards and the prevalence of bacteraemia

Data was collected from 27 wards which were categorized by the hospital into four main groups based on the type of patients; (i) Paediatric (ii) Intensive Care Units (ICUs) (iii) Surgical and (iv) Oncological. The study identified a higher prevalence of bacteraemia in the Oncology wards (68.4%) followed by Intensive Care Units (57.9%) (Table 02).

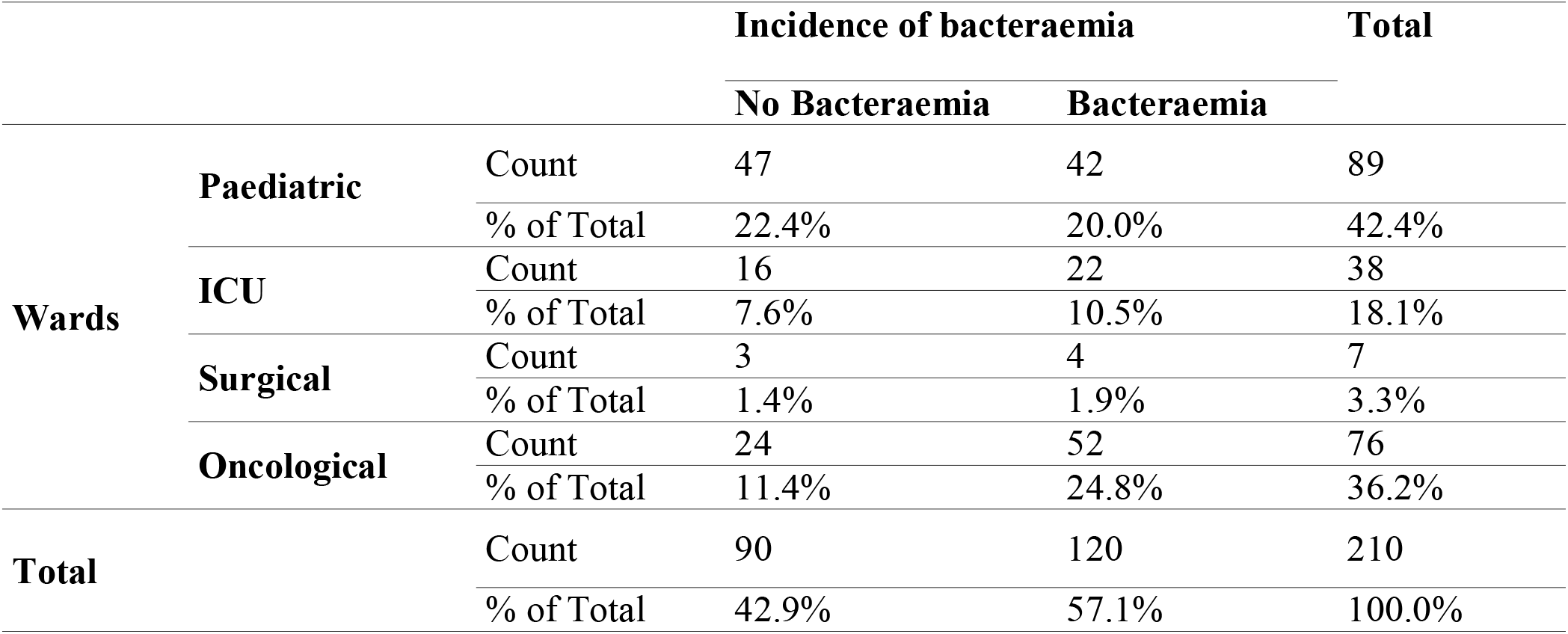

### Spectrum of Causative Organisms

Out of the 120 positive bacteraemic incidences identified, 31 (25.8%) were caused by lactose fermenting coliforms (Coliforms LF) and *Staphylococcus aureus* following up as the second most highly isolated pathogen attributing for 16 incidences (13.3%). Overall, Gram-negative pathogens accounted for the majority of the infections (49.17%) (Figure 01).

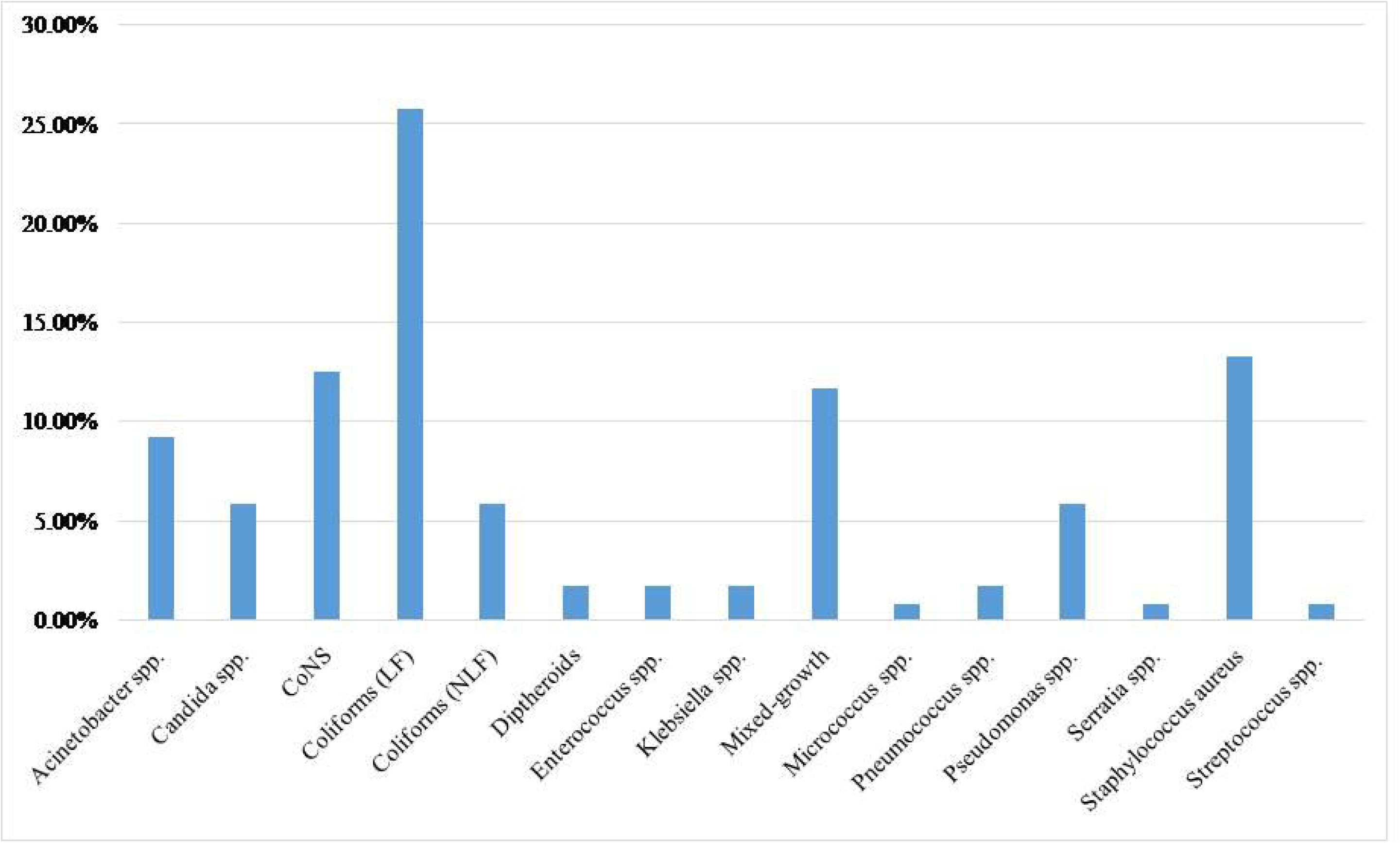

### The Association between Intravascular Devices and Bacteraemia

Several studies describe, normal flora at the insertion site or contamination of the catheter through direct contact during placement, and contamination of infusate as methods in which organisms could enter the blood stream of a patient with IVDs. 26.2% (55/210) of the total population were revealed to have IVDs, with 56.4% (31/55) of the IVD users being positive to bacteremia. Thus, the presence of an IVD was not a statistically significant factor contributing to the development of bacteraemia (*P*= 0.892).

### Spectrum of Microorganisms in Intravascular Devices (IVDs)

Multidrug-resistant (MDR) *Acinetobacter* specices were isolated from 29.0% (9/31) of the population studied with IVDs (Figure 05). MDR *Acinetobacter* species is defined as organisms displaying resistance to at least three classes of antibiotics including penicillins, cephalosporins, fluoroquinolones and aminoglycosides [15]. It is noteworthy that all *Acinetobacter* infections were isolated from patients with IVDs which means the organism was only prevalent among IVD users. Furthermore, *Acinetobacter* species displayed the highest resistance for antibiotics with sensitivity only to Polymixin B (Sensitivity of 93.3%). *Candida* species was reported as the second predominant organism isolated from IVDs (5/31 incidences, 16.1%) (Figure 02).

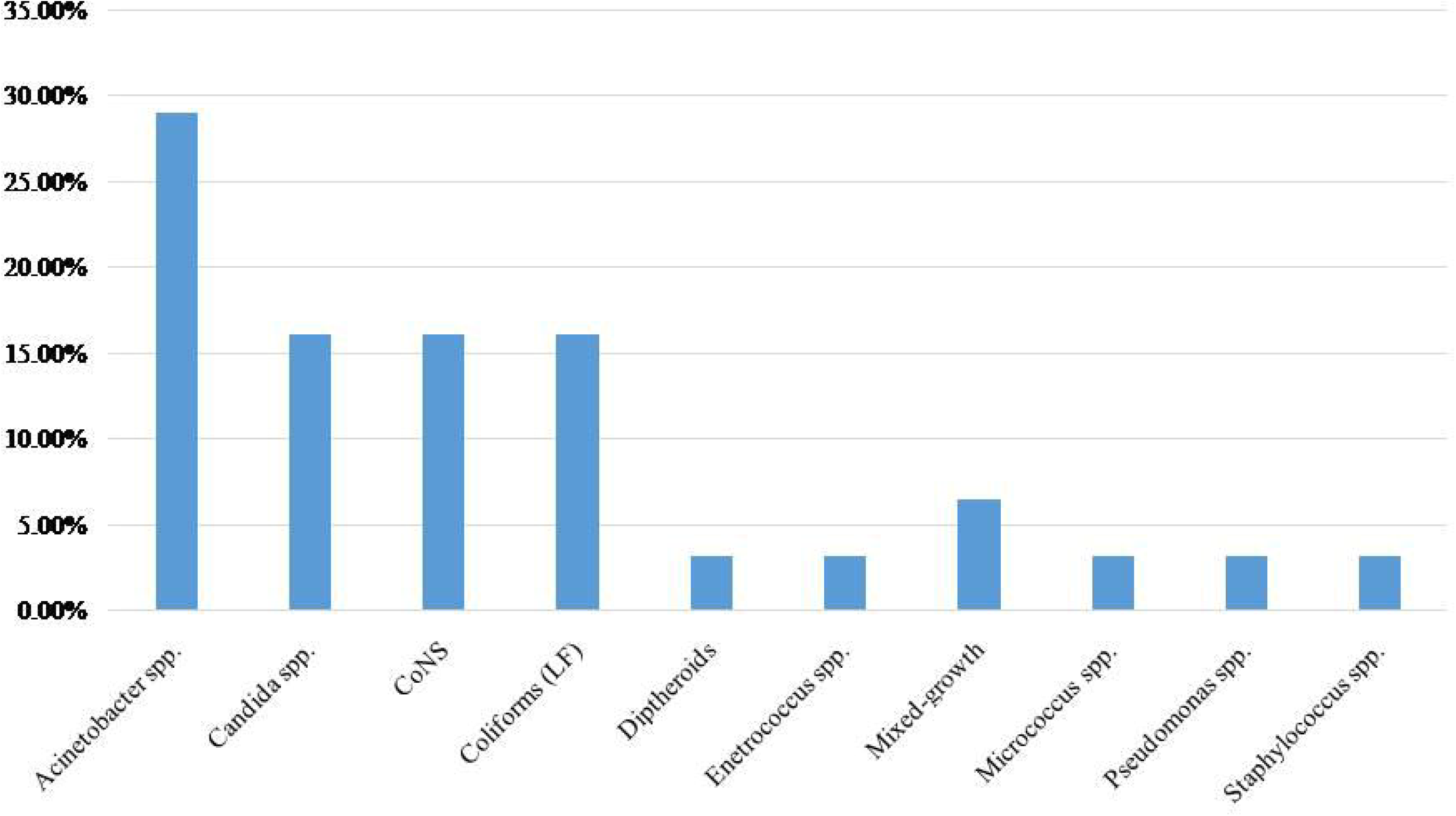

Our study reported only 8 Catheter Related Blood Stream Infections (CRBSIs) from 120 positive cases (6.7%) and majority of CRBSIs (3/8, 37.5%) were caused by *Acinetobacter* species.

### The Association between Neutropenia and Bacteraemia

Cancer patients are inevitably exposed to chemotherapy, corticosteroid drugs, stem cell transplantations and radiotherapy as part of their treatment regimen which result in neutropenia [5]. The current study identified majority of the cancer population to be non neutropenic (154/210, 73.3%) with a minority reporting positive to BSIs. Only 56 neutropenic incidences were reported of which 24.2% had bacteraemia while majority of the non neutropenic population had BSIs (59.1%). It was identified that neutropenia was not statistically significant in causing bacteraemia among cancer patients (*P*= 0.344) at this cancer hospital in South Asia.

### The Associations between Length of Hospital Stay and Bacteraemia

The Length of Stay (LOS) of cancer patients are affected by demographic factors, type of malignancy, the treatment regimen and infections caused by antibiotic resistant organisms [16,17]. On average, 3 samples were provided by a single patient, with hospital stay exceeding 3 days, over a period of two months. The present study identified LOS as a statistically significant factor contributing to the development of bacteraemia (*P*= 0.029, 95% CI), which supports the findings of the studies conducted previously (Figure 03). The average length of hospital stay was 14.1 days with 88/120 bacteremic incidences having an LOS greater than 3 days.

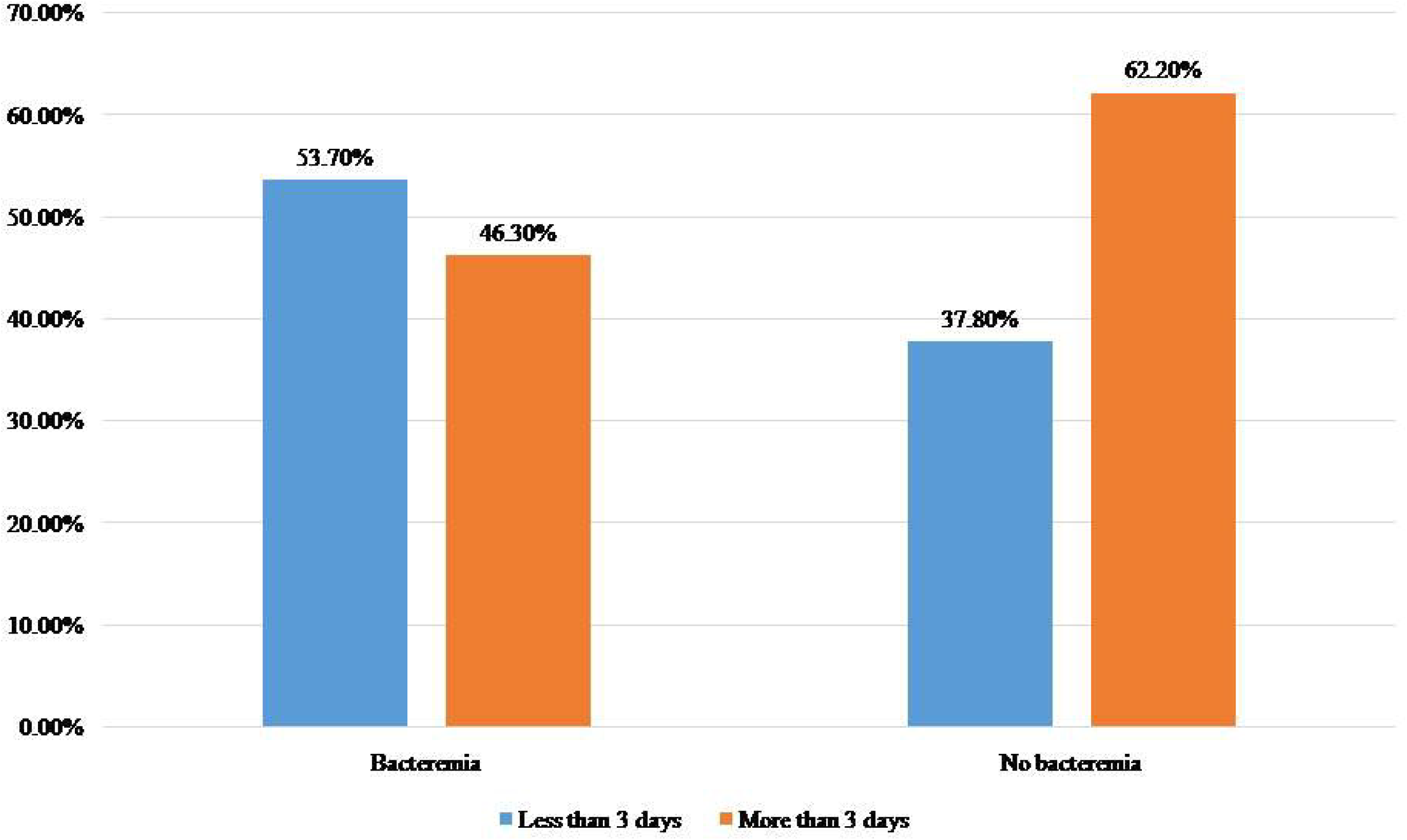

## Discussion

Our findings suggest that a Length of Stay (LOS) exceeding 3 days is a statistically significant factor contributing to the development of bacteremia among cancer patients at this South Asian Tertiary Cancer Centre, while individuals aged more than 60 years were identified as a vulnerable group.

Several studies define blood stream infections that give positive results following 48 to 72 hours of hospital admittance, as hospital-acquired bacteremia [13,18]. Thus, incidences with positive BSIs after a LOS of 3 days were identified as hospital-acquired blood stream infections (BSIs). Accordingly, we could deduce that 74.2% (89/120) of all bacteremic incidences of the present study were acquired within the hospital setting with other confounding factors such as invasive procedures, catheterization and age also coming into play. Studies by Welliver and McLaughlin [19] and Gardner and Carles [20] identified a higher prevalence of hospital acquired bacteremia among pediatric populations as opposed to adults. This was supported by our study, with majority (35.9%, 32/89) of hospital-acquired BSIs being identified among pediatric patients. A noteworthy observation was, identifying 75% (6/8) of Catheter Related Blood Stream Infections (CRBSIs) during prolonged hospital stay.

A meta-analytical study at John Hopkins University had revealed BSIs as the third leading hospital acquired infection [21] while data from National Nosocomial Surveillance Systems (NNIS) through January 1992 to June 2004 revealed a median rate of 1.8 to 5.2 CRBSIs per 1000 catheter days in Intensive Care Units (ICUs) in the US [22]. Similarly a study in the UK had identified CRBSIs contribute to 10% to 20% of the hospital acquired infections associated with increased ICU stays [23] which further indicates the global impact of hospital-acquired BSIs.

Majority of the incidences with a LOS exceeding 3 days were observed in patients with Acute Lymphoid Leukaemia (ALL) and Acute Myeloid Leukaemia (AML) (68.6% and 72.9% respectively). Several published literature reflected similar findings with a 3 fold greater risk of extended LOS from patients with AML [16,17]. Meanwhile, Kumari, Mishra and Mohan [9] revealed ALL patients to have the highest hospital duration. This indicates that patients with AML and ALL are at a greater risk in acquiring BSIs due to the lengthy hospital stays associated with the underlying malignancies, thus increased sterility measures and more medical attention should be given to such patients.

Patients above 60 years of age were revealed to be at a greater risk in acquiring BSIs which was in accordance with that of other studies. Nielsen [24] observed that elderly patients aged more than 65 years were at a greater risk in developing BSIs while Lenz et al. [25] and Al□Rawajfah, Stetzer and Beauchamp Hewitt [26] also reported similar findings in their studies which identifying elderly patients as a risk group in developing bacteraemia.

Gram-negative organisms were identified as the most predominant causative organism of BSIs (49.17%) while *Acinetobacter* was the most common species isolated from IVDs which was also observed by Chanock and Pizzo [27] and Aktaş et al [28]. In their study, Fukuta et al. [29] stated that the underlying malignancy does not contribute to infections by Multidrug-resistant (MDR) *Acinetobacter baumanni*, but rather acquired as a nosocomial infection. Kim et al. [30] describes a similar outcome and reveals that longer stays in the ICUs, increased the risk of acquiring MDR *A. baumanni* among cancer patients. Accordingly, our study identified all *Acinetobacter* related BSIs inpatients with a hospital stay exceeding 3 days. Candida was the second most common isolate identified with a higher prevalence among non-neutropenic patients. Candidemia is less prevalent among neutropenic populations as indicated in previous studies [31,32] with a greater incidence among patients with solid tumours. Rolston [33] has described the reason for this as the increased exposure of patients with solid tumours, to medical interventions such as catheterization and surgical procedures. Nonetheless, our study did not show a difference between patients with haematological malignancies and solid tumours since each reported two cases of candidaemia. According to Walsh and Rex [34] candidaemia is ranked as the 3^rd^ to 4^th^ most common nosocomial BSI. This becomes factual with the present study where 3 out of the 5 candidaemic incidences were in patients with a Length of Stay (LOS) exceeding 3 days which means the infections are acquired from the hospital setting.

An interesting finding was the higher prevalence of BSIs among the non-neutropenic population, which is an indication that neutropenia is not a risk factor in causing bacteraemia in the present population. This contradicted several studies which revealed neutropenia and the degree of neutropenia as risk factors for bacteremia [35,36]. Our study recorded 136 haematological malignancies of which 52 (57.1%) were non-neutropenic however, the neutrophil function was not assessed thus, the qualitative nature of neutrophils and its affect in increasing the risk of BSIs could not be determined.

Our study also revealed that BSIs were less prevalent among the patients with an IVD which also contradicted the findings of many studies which considered IVDs as a risk factor for BSIs [37–39]. This could be a result of complications such as surgeries, immunosuppression and the underlying malignancies itself which would result in bacteraemia, independent of the presence of an IVD.

In conclusion, a hospital stay exceeding three days was identified as a risk factor contributing to the development of bacteraemia while neutropenia and the presence of an intravascular device were identified to have no impact on the risk in causing BSIs in the present study. Therefore, patients who are at a risk of prolonged hospital stays, such as patients diagnosed with AML and ALL and the paediatric population, should be given additional medical attention ensuring sterility during medical procedures. Furthermore, bacteraemia was more prevalent in old age, especially among patients exceeding 60 years, which was identified as a vulnerable category, therefore hospital staff should be more vigilant when dealing with the senile population. Thus, through this study it is evident that hospital-acquired BSIs imposes an additional burden on cancer patients, therefore it is important that hospitals take necessary measures to minimize its occurrence.

